# O-GlcNAc glycosylation orchestrates fate decision and niche function of bone marrow stromal progenitors

**DOI:** 10.1101/2022.12.21.521379

**Authors:** Zengdi Zhang, Zan Huang, Mohamed Awad, Mohammed Elsalanty, James Cray, Lauren E. Ball, Jason C. Maynard, Alma L. Burlingame, Hu Zeng, Kim C. Mansky, Hai-Bin Ruan

## Abstract

In mammals, interactions between the bone marrow (BM) stroma and hematopoietic progenitors contribute to bone-BM homeostasis. Perinatal bone growth and ossification provide a microenvironment for the transition to definitive hematopoiesis; however, mechanisms and interactions orchestrating the development of skeletal and hematopoietic systems remain largely unknown. Here, we establish intracellular O-linked β-N-acetylglucosamine (O-GlcNAc) modification as a posttranslational switch that dictates the differentiation fate and niche function of early BM stromal cells (BMSCs). By modifying and activating RUNX2, O-GlcNAcylation promotes osteogenic differentiation of BMSCs and stromal IL-7 expression to support lymphopoiesis. In contrast, C/EBPβ-dependent marrow adipogenesis and expression of myelopoietic stem cell factor (SCF) is inhibited by O-GlcNAcylation. Ablating O-GlcNAc transferase (OGT) in BMSCs leads to impaired bone formation, increased marrow adiposity, as well as defective B-cell lymphopoiesis and myeloid overproduction in mice. Thus, the balance of osteogenic and adipogenic differentiation of BMSCs is determined by reciprocal O-GlcNAc regulation of transcription factors, which simultaneously shapes the hematopoietic niche.

## INTRODUCTION

Mammalian bones support body structure, protect vital organs, and allow body movement. In addition, they provide an environment for hematopoiesis in the bone marrow (BM). Most bones in mammals are formed through endochondral ossification, which is initiated by mesenchymal condensation, followed by the differentiation of chondrocytes and perichondrial progenitors (Kobayashi and Kronenberg, 2021). Perichondrial progenitors expressing Osterix (Osx) co-migrate with blood vessels into the primary ossification center, giving rise to osteoblasts and transient stromal cells in the nascent BM cavity (Chen et al., 2014; Liu et al., 2013; Maes et al., 2010; Mizoguchi et al., 2014). At the perinatal stage, Osx^+^ progenitors contribute to osteolineages and long-lived BM stromal cells (BMSCs) that exhibit trilineage differentiation potential into osteocytes, chondrocytes, and adipocytes.

The decision of BMSC fate is controlled by a transcriptional network of pro-osteogenic and anti-adipogenic transcription factors that pre-establishes osteogenic enhancers in BMSCs for rapid bone formation (Rauch et al., 2019). RUNX family transcription factor 2 (RUNX2), by regulating osteogenic genes including *Osx*, determines the osteoblast lineage from the multipotent BMSCs. Mice with *Runx2* mutations completely lack skeletal ossification and die of respiratory failure (Komori et al., 1997). *Runx2*-haploinsufficient mice show specific skeletal abnormalities characteristic of human cleidocranial dysplasia (CCD), including persistent fontanels, delayed closure of cranial sutures, rudimentary clavicles, and dental abnormalities (Otto et al., 1997; Takarada et al., 2016). On the other hand, adipogenesis is driven by downregulation of pro-osteogenic factors, remodeling of the chromatin, and activation of adipogenic transcription factors, such as C/EBPs and PPARγ (Rauch *et al*., 2019). BM adiposity is associated with bone loss in osteoporosis caused by aging, menopause, and anorexia nervosa (Bethel et al., 2013; Fazeli et al., 2013; Liu et al., 2015; Scheller et al., 2016). However, it is incompletely understood how these distinct types of transcription factors act cooperatively to determine lineage differentiation during neonatal skeletal development.

BMSCs and their lineage-differentiated progeny (e.g., osteoblasts and adipocytes) provide a niche microenvironment for hematopoiesis (Bianco and Robey, 2015; Calvi and Link, 2015; Morrison and Scadden, 2014; Wei and Frenette, 2018). Recent studies using single cell technologies and lineage tracing experiments have started to unveil the complexity and heterogeneity of niche cell types, niche factors, and their actions. For example, BMSC-derived stem cell factor (SCF, encoded by the *Kitl* gene) and CXC chemokine ligand 12 (CXCL12) are required for the maintenance and differentiation of hematopoietic stem/progenitor cells (HSPCs) (Asada et al., 2017; Ding et al., 2012). A prominent subpopulation of perivascular BMSCs express adipocyte markers (Dolgalev and Tikhonova, 2021; Zhong et al., 2020; Zhou et al., 2017), and support steady-state and metabolic-stressed myelopoiesis by secreting SCF (Zhang et al., 2019). Meanwhile, osteolineage cells are crucial for lymphopoiesis (Wei and Frenette, 2018). Depleting *Osx*^+^ cells halts B cell maturation and causes immune failure (Yu et al., 2016). IL-7, the most crucial factor for lymphoid progenitors, is expressed by a subset of BMSCs (Fistonich et al., 2018). While it is well accepted that myeloid and lymphoid progenitors may reside in distinct BM niches, it is unclear how BMSC heterogeneity is established during early development and whether cytokine expression is coordinated and controlled by the fate-defining transcriptional network in BMSCs.

Post-translational modifications (PTMs), including phosphorylation, acetylation, and ubiquitination, allow the precise regulation of stability, localization, and activity of BM transcriptional factors, such as RUNX2 (Chen et al., 2021; Kim et al., 2020), C/EBPs (Wang et al., 2022), and PPARγ (Brunmeir and Xu, 2018). It remains poorly defined how these modifications are coordinated in a spatio-temporal manner to calibrate skeletal development. Thousands of intracellular proteins are dynamically modified by a single O-linked N-Acetylglucosamine (O-GlcNAc) moiety at serine or threonine residues, termed O-GlcNAcylation (Hart et al., 2007; Ruan et al., 2013b; Yang and Qian, 2017). O-GlcNAc transferase (OGT), using UDP-GlcNAc derived from the hexosamine biosynthetic pathway as the substrate, controls diverse biological processes such as gene transcription, protein stability, and cell signaling (Hanover et al., 2012; Ruan et al., 2014; Ruan et al., 2012; Ruan et al., 2013a). In cell culture, O-GlcNAcylation promotes osteogenesis (Kim et al., 2007; Nagel and Ball, 2014) and suppresses adipogenesis (Ji et al., 2012). However, the physiological relevance of O-GlcNAcylation in skeletal development and remodeling has not been established. Here, we studied OGT in balancing osteogenic versus adipogenic programs and in controlling niche function of BMSC in mice. The multifaceted role of protein O-GlcNAcylation is achieved through reciprocal regulation of pro-osteogenic, pro-lymphopoietic RUNX2 and pro-adipogenic, pro-myelopoietic C/EBPβ.

## RESULTS

### Loss of OGT in perinatal BMSCs leads to bone loss and marrow adiposity

To determine the in vivo role of protein O-GlcNAcylation in bone development, we deleted the X Chromosome-located *Ogt* gene using the *Osx-GFP:Cre* mice (**Figure 1A**). Compared to *Osx-Cre^+^* only littermate controls, newborn *Osx*^Δ*Ogt*^ mice showed no obvious change in long bone formation (**Figure 1B**) but had a profound defect in the mineralization of flat bones of the calvaria (**Figure 1C**), suggesting impaired intramembranous ossification during the prenatal stage.

**Figure 1.**
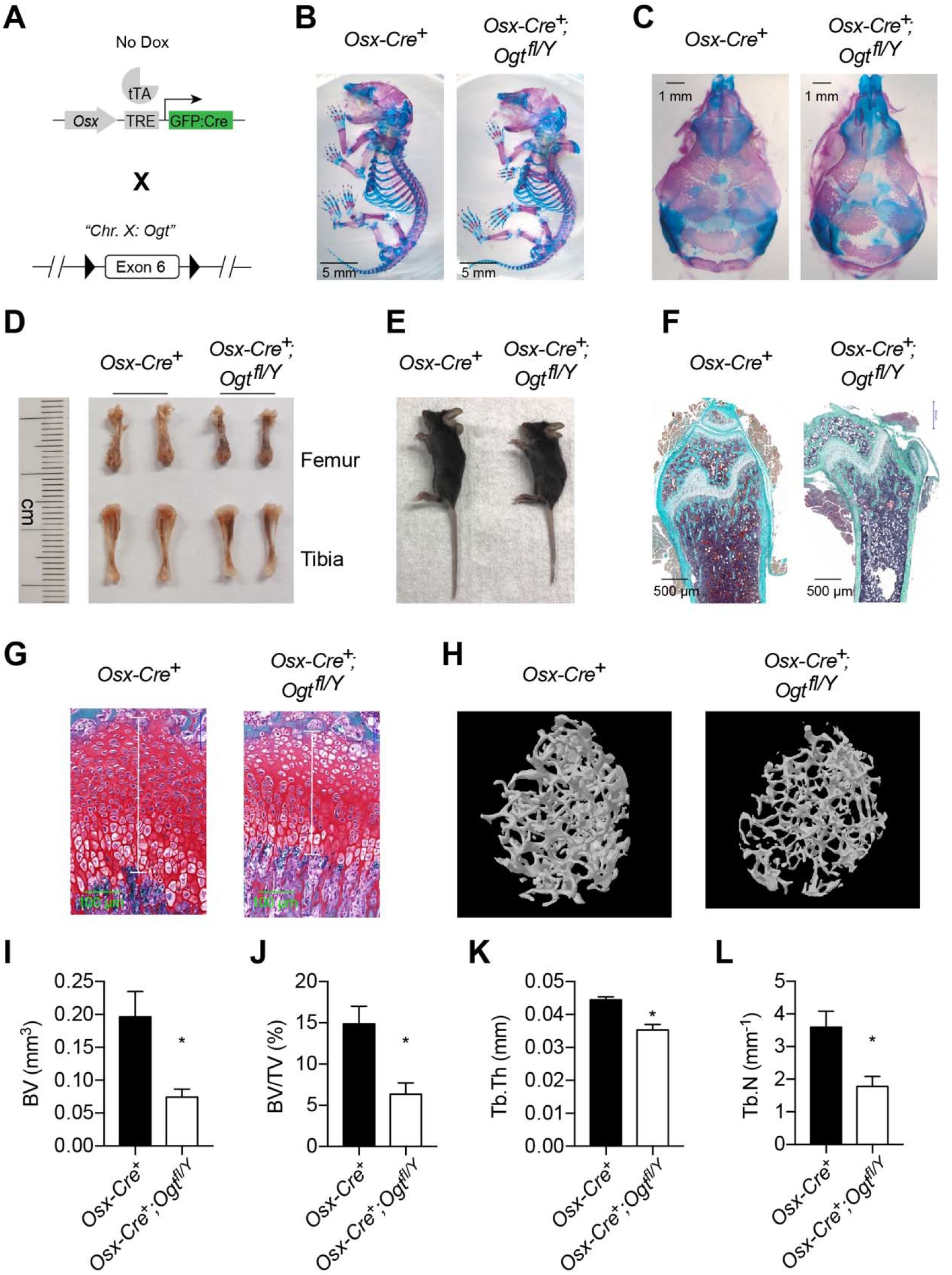
Impaired osteogenesis in *Osx*^Δ*Ogt*^ mice. (**A**) Mating strategy to generate *Osx*^Δ*Ogt*^ mice. Note that the *Ogt* gene is located on Chr. X, thus males are hemizygous *Ogt^fl/Y^*. (**B, C**) Whole mount Alizarin red and Alcian blue staining of newborn mice. (**D, E**) Long bone length (D) and gross morphology of 4-6 weeks old mice. (**F-G**) Goldner’s trichrome (F) and Safranin O (G) staining of femurs from 4-week-old mice. (**H-L**) Micro-CT of 6-week-old mice (H, n = 3-4). Bone volume (BV, I), BV/tissue volume ratio (BV/TV, J), trabecular thickness (Tb.Th, K), and trabecular number (Tb.N, L) were calculated. Data are presented as mean ± SEM. *, p < 0.05 by unpaired student’s *t*-test.

At 4-6 weeks of age, *Osx*^Δ*Ogt*^ mice were modestly shorter than the controls (**Figure 1D, E**). Histological analyses showed decreased bone volume and osteoblast number (**Figure 1F**) and shortened growth plate (**Figure 1G**) in *Osx*^Δ*Ogt*^ mice. Micro-CT scanning further showed that *Osx*^Δ*Ogt*^ mice had reduced trabecular bone volume, bone volume to tissue volume ratio, trabecular thickness, and trabecular numbers in the distal femur (**Figure 1H-L**). *Osx*^Δ*Ogt*^ mice represent typical bone and dental defects (**Figure 1—figure supplement 1**) as observed in *Runx2*-haploinsufficient mice (Otto *et al*., 1997; Takarada *et al*., 2016), suggesting that O-GlcNAcylation might control RUNX2 function.

### RUNX2 O-GlcNAcylation promotes osteogenesis

To investigate how OGT controls osteogenic differentiation of BMSCs, we first isolated primary BMSCs from control and *Osx*^Δ*Ogt*^ mice and induced them into osteoblast cells. Alkaline phosphatase staining revealed a reduction in mineralization of *Osx*^Δ*Ogt*^ BMSCs **(Figure 2A**). Similarly, treating mesenchymal C3H10T1/2 cells with an OGT inhibitor, OSMI-1, reduced mineralization (**Figure 2B**) and ablated calcium deposition (**Figure 2C**) after osteogenic differentiation. Parathyroid hormone (PTH) is a bone anabolic agent that requires RUNX2-dependent signaling (Krishnan et al., 2003). We found that PTH treatment of C3H10T1/2 cells increased global protein O-GlcNAcylation (**Figure 2D**). The ability of PTH to activate osteogenesis is completely abolished when OGT was inhibited by OSMI-1 (**Figure 2E**). Pharmacological activation of O-GlcNAcylation enhances RUNX2 activity and promotes osteogenic differentiation (Kim *et al*., 2007; Nagel and Ball, 2014). We mutated three known O-GlcNAc sites on RUNX2, Ser 32 and Ser 33 in the N-terminal transactivation domain and Ser 371 in the proline/serine/threonine-rich domain (**Figure 2F**), to alanine (3A), and found mutant RUNX2 possessed less O-GlcNAcylation (**Figure 2G**). O-GlcNAcase (OGA) inhibition by Thiamet-G (TMG) increased O-GlcNAcylation of wildtype (WT) RUNX2, but to a much less extent in the 3A mutant (**Figure 2G**). OGT inhibition by OSMI-1 or O-GlcNAc mutation both impaired the transcriptional activity of RUNX2 on a luciferase reporter (**Figure 2H**). OSMI-1 could still suppress of luciferase activity of RUNX2-3A (**Figure 2H**), suggesting additional, unidentified O-GlcNAc sites (**Figure 2G**), which requires future investigation. Nevertheless, when overexpressed in C3H10T1/2 cells, RUNX2-3A substantially lost the ability to induce osteogenic differentiation (**Figure 2I**) or RUNX2-target gene expression (**Figure 2J**). These data demonstrate that O-GlcNAcylation is essential for RUNX2 activity and osteogenesis.

**Figure 2.**
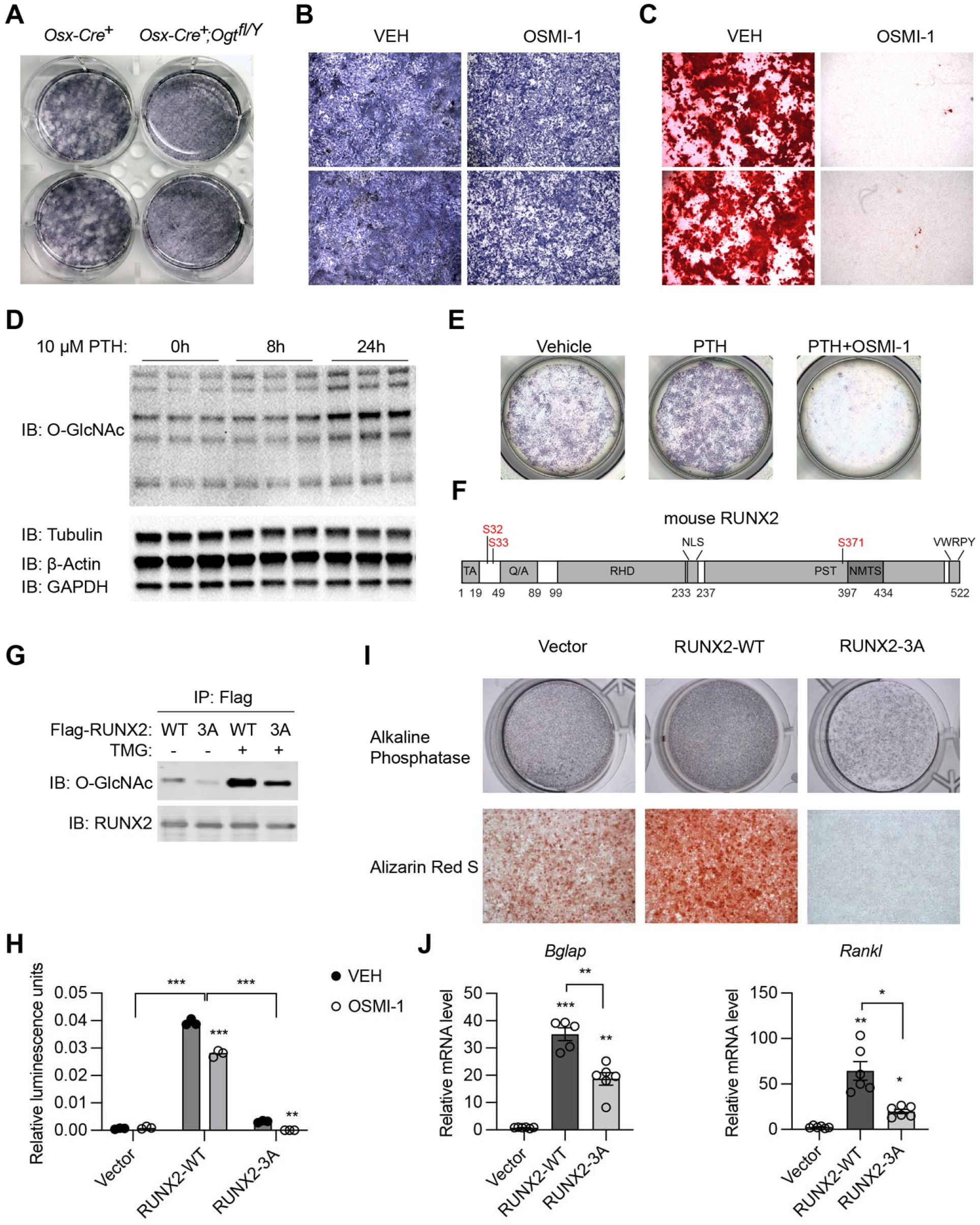
RUNX2 O-GlcNAcylation is required for osteogenesis. (**A**) Alkaline phosphatase (ALP) staining of control and *Osx*^Δ*Ogt*^ BMSCs differentiated to the osteogenic lineage. (**B, C**) Primary BMSCs, in the presence or absence of the OGT inhibitor TMG, were induced for osteogenesis and stained for ALP (B) and Alizarin Red S (C). (D) Primary BMSCs were treated with PTH for the indicated time and subjected to Western blotting of total protein O-GlcNAcylation. (**E**) BMSCs were treated with PTH alone or together with TMG, osteogenic differentiated, and stained for ALP (n = 3). (**G**) Flag-tagged wildtype (WT) and O-GlcNAc mutant (3A) RUNX2 plasmids were overexpressed in HEK293 cells, and their O-GlcNAcylation was determined by Flag immunoprecipitation followed with O-GlcNAc Western blot. (**H**) 6xOSE-luciferase activity in COS-7 cells transfected with WT- or 3A-mutant RUNX2, in the presence or absence of the OGT inhibitor, OSMI-1. (**I, J**) C3H10T1/2 cells with lentiviral overexpression of RUNX2 were osteogenically differentiated and stained with ALP or Alizarin Red S (I). Expression of *Bglap* and *Rankl* was determined by RT-qPCR (J). Data are presented as mean ± SEM. *, p < 0.05; **, p < 0.01; and ***, p < 0.001 by one-way ANOVA (J). Representative images from at least 3 biological replicates were shown in A, B, C, E, and I.

In adult mice, *Osx* expression is restricted to osteoblast precursors. We treated *Osx*^Δ*Ogt*^ mice from pregnancy with doxycycline (Dox) and withdrew Dox at 10 weeks of age to induce Cre expression and OGT depletion only during adulthood (**Figure 3A**). Micro-CT showed that *Osx*^Δ*Ogt*^ mice had reduced bone volume, trabecular thickness, and bone mineral density (**Figure 3B-E**). Together, these results support the functional indispensability of OGT in the committed osteolineage for adult trabecular bone remodeling.

**Figure 3.**
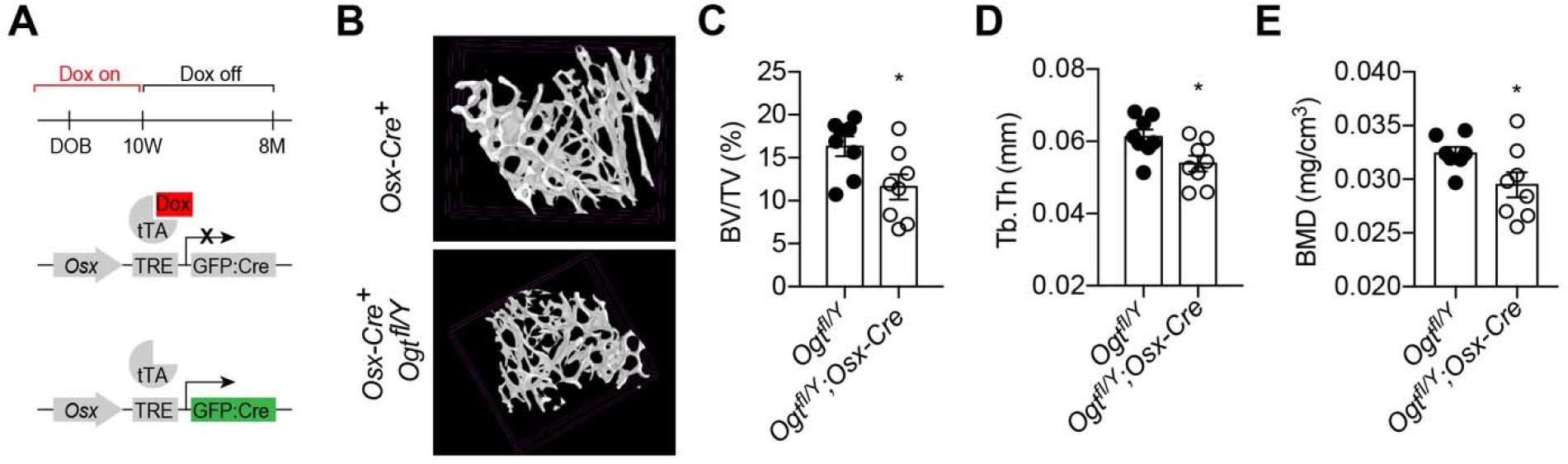
Adult-onset deletion of OGT impairs trabecular bone formation. (**A**) Dox treatment timeline in *Osx*^Δ*Ogt*^ to achieve osteoblast-specific deletion of OGT. (**B-D**) Micro-CT (B) showing reduced bone volume/tissue volume (C), trabecular thickness (D), and trabecular bone mineral density (E). Data are presented as mean ± SEM.*, p < 0.05 by unpaired student’s t-test.

### C/EBPβ O-GlcNAcylation inhibits the adipogenic specification of BMSCs

The osteogenic and adipogenic differentiation of BMSCs is generally considered mutually exclusive (Ambrosi et al., 2017). Concomitant with bone loss, we observed a massive accumulation of adipocytes in the bone marrow of *Osx*^Δ*Ogt*^ mice, shown by hematoxylin & eosin staining (**Figure 4A**) and immune-staining of the lipid droplet protein - perilipin (**Figure 4B**). Flow cytometric analysis of BMSCs revealed that *Osx*^Δ*Ogt*^ mice possessed more PDGFRα^+^/VCAM1^+^ adipogenic progenitors than littermate controls (**Figure 4C**). To directly test if OGT deficiency biases BMSC differentiation toward the adipogenic lineage, we first induced the adipogenic differentiation of primary BMSCs and found increased lipid deposition in *Osx*^Δ*Ogt*^ mice (**Figure 4D**). Even under an osteogenic induction condition, adipo-lineage markers such as *Adipoq* and *Vcam1* were significantly upregulated by OGT deficiency (**Figure 4E, F**). Furthermore, treating C3H10T1/2 mesenchymal cells with an OGA inhibitor TMG to increase protein O-GlcNAcylation, was able to substantially reduce perilipin protein expression (**Figure 4G**). These data indicate that OGT inhibits the adipogenic program of BMSCs.

**Figure 4.**
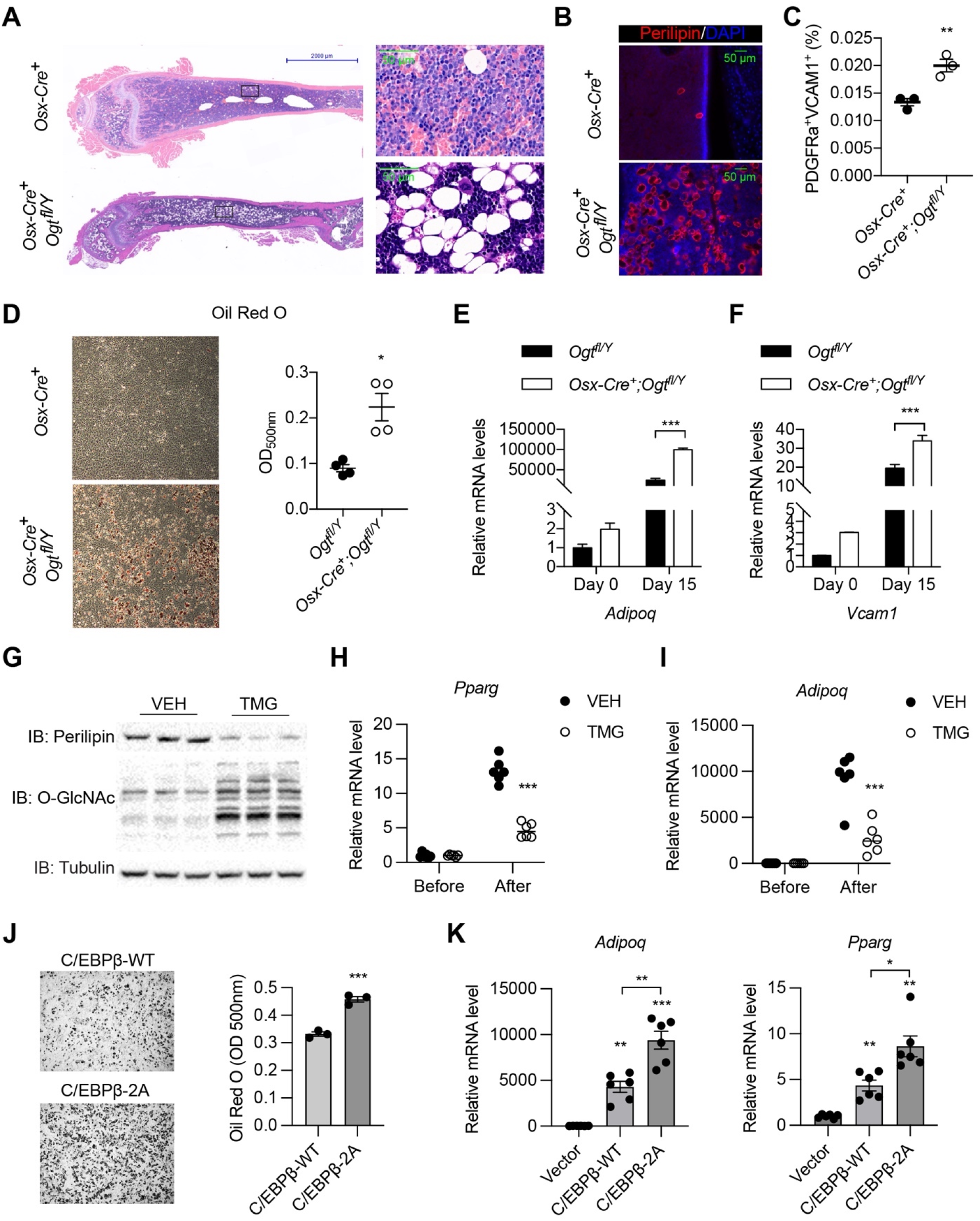
O-GlcNAcylation inhibits BM adipogenesis. (**A, B**) H&E (A) and Perilipin immunofluorescent staining (B) on femur sections from 4-week-old mice. (**C**) Flow cytometric quantification of PDGFRa^+^VCAM1^+^ preadipocytes frequencies within the live BM cells (n = 3). (**D**) Adipogenic differentiation of primary BMSCs from control and *Osx*^Δ*Ogt*^ mice. Lipid was stained with Oil Red O and quantified to the right (n = 4). (**E, F**) Primary BMSCs were osteogenic differentiated for 0 or 15 days. Expression of *Adipoq* (E) and *Vcam1* (F) genes was determined by RT-qPCR (n = 3). (**G-I**) C3H10T1/2 cells, treated with or without TMG, were adipogenic differentiated. Western blotting for perilipin and O-GlcNAc of differentiated cells (G) and RT-qPCR for adipogenic marker *Pparg* (H) and *Adipoq* (I) expression. (**J, K**) Adipogenic differentiation of C3H10T1/2 cells infected with lentiviral C/EBPβ. Oil Red O was stained and quantified (J). *Pparg* and *Adipoq* gene expression was determined by RT-qPCR (K). Data are presented as mean ± SEM.*, p < 0.05; **, p < 0.01; and ***, p < 0.001 by unpaired student’s t-test (C, J), one-way ANOVA (K), and two-way ANOVA (E, F, H, I).

We went on to determine the O-GlcNAc targets of OGT in suppressing adipogenesis. As an osteogenic regulator, RUNX2 also reciprocally suppresses the adipogenic program (Ahrends et al., 2014). However, such suppression was not dependent on O-GlcNAcylation, because O-GlcNAc-deficient RUNX2 displayed similar efficiency as the wildtype protein to reduce lipid deposition and perilipin expression in differentiated C3H10T1/2 cells (**Figure 4—figure supplement 1**). PPARγ1 is O-GlcNAcylated at T54 in the A/B activation domain (Ji *et al*., 2012), corresponding to T84 in PPARγ2, the major isoform in adipocytes (**Figure 4—figure supplement 2A**). Mutating T84 in PPARγ2 did not ablate the ability of the OGA inhibitor TMG to suppress adipogenesis in C3H10T1/2 cells (data not shown), suggesting the existence of other unidentified O-GlcNAc sites on PPARγ2 or other target proteins than PPARγ2. Through mass spectrometry, we were able to map four additional O-GlcNAc sites on PPARγ2 (**Figure 4— figure supplement 2A** and **source data 1**). Intriguingly, mutating these 4 sites or together with T84 to alanine, render PPARγ2 incompetent to induce transcription and adipogenesis (**Figure 4—figure supplement 2B, C**). It suggests that PPARγ2 O-GlcNAcylation is essential for adipocyte maturation, but likely does not mediate the anti-adipogenic effect of OGT in perinatal BMSCs.

We then looked to C/EBPβ, an early transcription factor that specifies the adipogenic fate of BMSCs (Cao et al., 1991; Darlington et al., 1998). It has been reported that OGT modifies C/EBPβ to inhibit its transcriptional activity (Li et al., 2009; Qian et al., 2018). As expected, ablating O-GlcNAcylation of C/EBPβ (2A mutation) promotes adipogenic differentiation of C3H10T1/2 cells (**Figure 4H, I**). Taken together, we concluded that, by O-GlcNAcylating and reciprocally regulating RUNX2 and C/EBPβ, OGT is required for the proper allocation of skeletal progenitors into osteogenic versus adipogenic lineages during development.

### OGT deficiency disrupts the BM niche

Skeletal development is concomitant with the establishment of definitive hematopoiesis in the BM. To test if OGT deficiency affects the niche function of *Osx1*^+^ cells for B-cell lymphopoiesis, we performed flow cytometry analyses of bone marrow of 4-week-old mice (**Figure 5A, figure supplement 1, and source data 1**) (Hardy et al., 1991). No changes in the percentage of lineage^-^Sca-1^+^Kit^+^ progenitor cells, common lymphoid progenitors (CLPs), Fraction A that contains pre-pro-B cells were observed between control and *Osx*^ΔOgt^ mice (**Figure 5B-D**). While frequencies of Fraction B and C pro-B, pre-B, and immature B in *Osx*^Δ*Ogt*^ mice were drastically reduced (**Figure 5E-I**), demonstrating a developmental blockage from pre-pro-B to pro-B cells. In the peripheral blood, there was specific loss of CD19^+^B220^+^ B cells but not CD4^+^ or CD8^+^ T cells (**Figure 5J-L**). B-cell dysfunction observed here was similar to the phenotype in mice when all Osx^+^ cells were depleted (Yu *et al*., 2016) or IL-7 was deleted in BMSCs (Cordeiro Gomes et al., 2016), indicating that O-GlcNAcylation is essential for the Osx^+^ lineage cells to establish a niche environment for B-cell lymphopoiesis.

**Figure 5.**
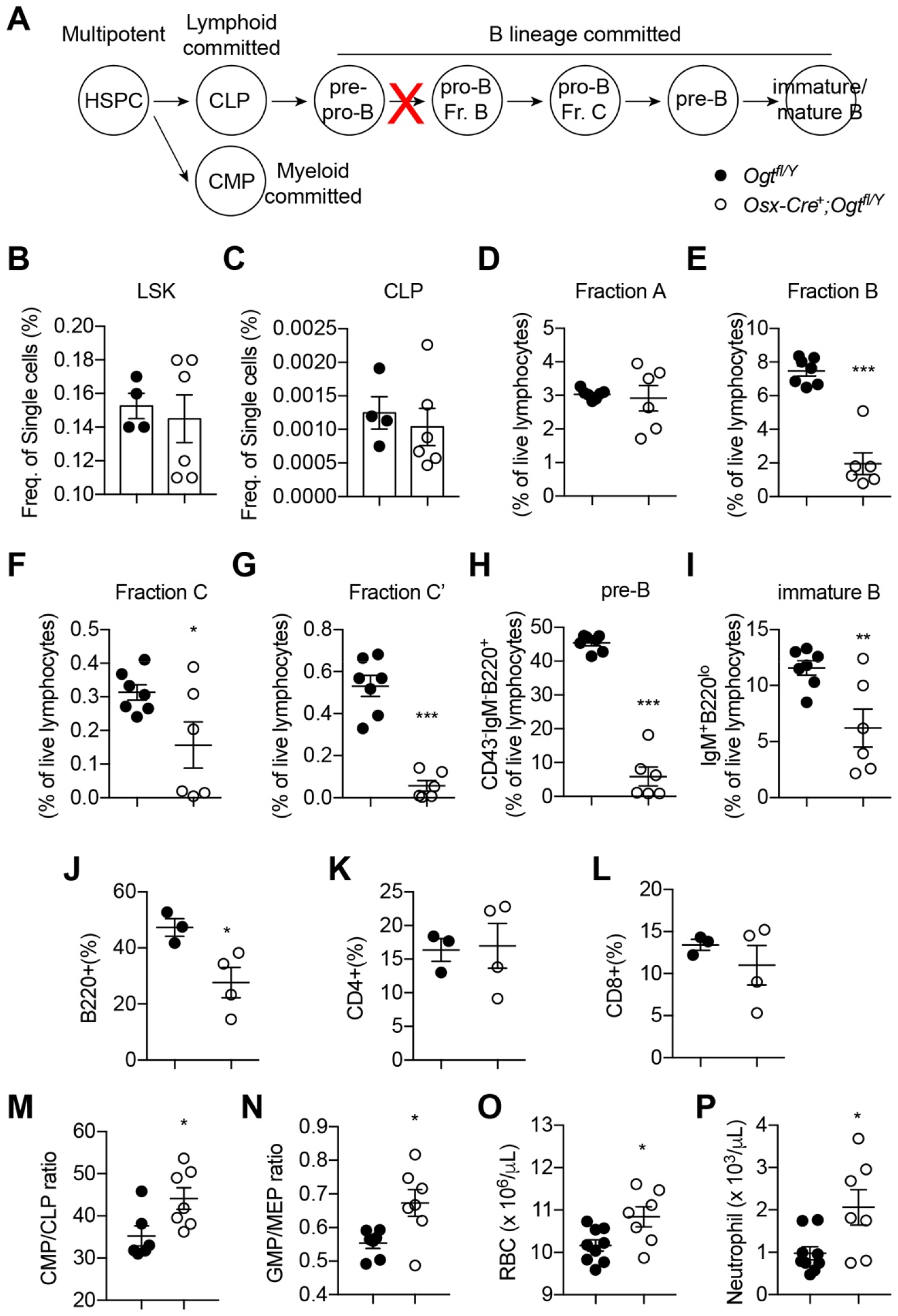
Impaired B lymphopoiesis and myeloid skewing in *Osx*^Δ*Ogt*^ mice. (**A**) Schematic view of B cell development in the BM and blockade by stromal OGT deficiency (red X). (**B-C**) Flow cytometric quantification of LSK (B) and CLP (C) among live BM cells (n = 4-6). (**D-I**) Flow cytometric quantification of fraction A (D), fraction B (E), fraction C (F), fraction C’ (G), fraction D (H), and immature B (I) frequencies among live BM lymphocytes (n = 6-7). (**J-L**) Flow cytometric quantification of B220^+^ B cell (J), CD4^+^ T cell (K), and CD8^+^ T cell (L) percentages in the blood (n = 3-4). (**M, N**) CMP/CLP ratio (M) and GMP/EMP ratio (N) in the BM (n = 6-7). (**O, P**) Complete blood counting showing numbers of RBC (O) and neutrophil (P) (n = 7-9). Data are presented as mean ± SEM.*, p < 0.05; **, p < 0.01; and ***, p < 0.001 by unpaired student’s t-test.

BM adiposity is associated with myeloid overproduction in conditions including aging, irradiation (Ho et al., 2019), osteopenia (Kajkenova et al., 1997), and obesity (Singer et al., 2014), indicating the supportive function of marrow adipocytes on demand-adapted myelopoiesis. Consistently with the increased BM adiposity found in *Osx*^Δ*Ogt*^ mice, we also observed biased HSPC differentiation toward the myeloid lineage, as shown by increased ratio of common myeloid progenitor (CMP) to common lymphoid progenitors (CLPs) and ratio of granulocyte-monocyte progenitors (GMP) to megakaryocyte-erythrocyte progenitors (MEP) in the BM (**Figure 5M, N**). As a result, increased numbers of red blood cells and neutrophils were observed in the blood of *Osx*^Δ*Ogt*^ mice (**Figure 5O, P**). Together, these results demonstrate that OGT deficiency in neonatal BMSCs establishes a BM environment that promotes myelopoiesis and simultaneously impairs B cell development.

### Transcriptional regulation of niche cytokines by RUNX2 and C/EBPβ O-GlcNAcylation

BMSC-derived SCF (encoded by the *Kitl* gene) and IL-7 are required for the myeloid differentiation and B-cell development, respectively (Asada *et al*., 2017; Cordeiro Gomes *et al*., 2016; Ding *et al*., 2012). We sought to test if their expression is controlled by the same transcriptional network determining BMSC fate. Adipogenic differentiation of mesenchymal C3H10T1/2 cells concomitantly increased *Kitl* while decreased *Il7* gene expression (**Figure 6A, B**). Simultaneous treatment with the OGT inhibitor OSMI-I dampened *Il7* expression before differentiation but enhanced *Kitl* expression in differentiated adipocytes (**Figure 6A, B**). On the other hand, osteogenic differentiation suppressed *Kitl* transcription, which could be further inhibited by TMG that elevated global O-GlcNAcylation (**Figure 6C**). While *Il7* mRNA levels were not evidently affected by osteogenic differentiation, TMG stimulated its expression (**Figure 6D**). O-GlcNAcylation inhibits the adipogenesis specified by C/EBPβ but supports osteogenesis determined by RUNX2. In concert, C/EBPβ overexpression in C3H10T1/2 cells activated *Kitl* transcription and suppressed *Il7* expression, which was further exacerbated by O-GlcNAc-deficient C/EBPβ (**Figure 6E, F**). However, RUNX2 overexpression decreased *Kitl* mRNA levels (**Figure 6G**). When compared to the wildtype, O-GlcNAc-defective RUNX2 was impaired in inducing *Il7* expression (**Figure 6H**). Collectively, these results reveal that protein O-GlcNAcylation, by acting on BMSC lineage transcriptional factors, establishes a pro-lymphopoietic niche during neonatal bone development and at the same time prevents the myeloid-skewing, adipogenic BM environment.

**Figure 6.**
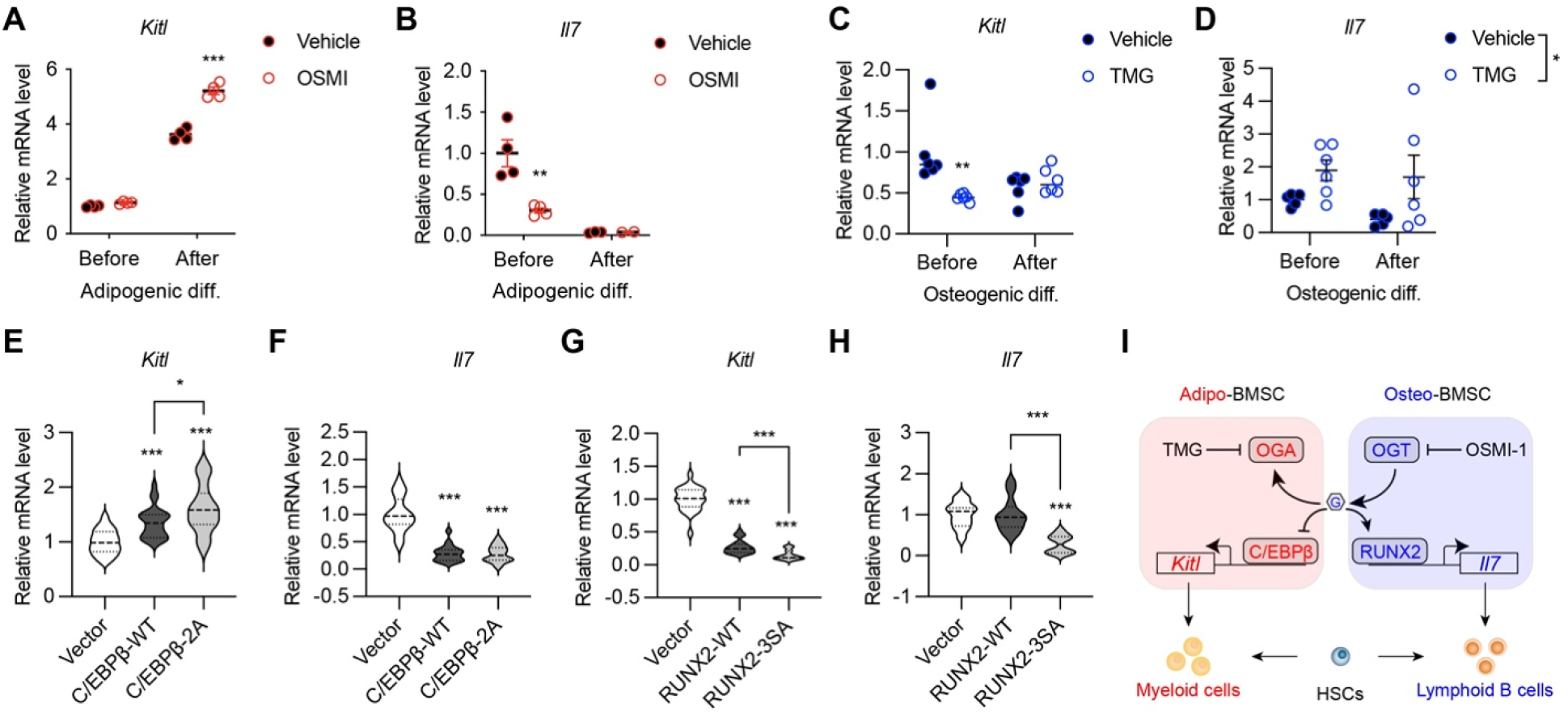
O-GlcNAc regulation of niche cytokine expression. (**A, B**) C3H10T1/2 cells were treated with vehicle or OGT inhibitor OSMI and differentiated for adipocytes (n = 4). *Kitl* (A) and *Il7* (B) gene expression was determined by RT-qPCR. (**C, D**) C3H10T1/2 cells were treated with vehicle or OGA inhibitor TMG and induced for osteogenic differentiation (n = 6). *Kitl* (C) and *Il7* (D) gene expression was determined by RT-qPCR. (**E-H**) C3H10T1/2 cells were infected with lentiviruses expressing WT and O-GlcNAc-deficient C/EBPβ (E, F) or RUNX2 (G, H). Expression *Kitl* (E, G) and *Il7* (F, H) was measured by RT-qPCR (n = 6). (**I**) Proposed action of protein O-GlcNAcylation in regulating the BMSC niche function. Data are presented as mean ± SEM.*, p < 0.05; **, p < 0.01; ***, p < 0.001 by two-way ANOVA (A-D) and one-way ANOVA (E-H).

## DISCUSSION

Post-translational modification networks exist in the bone-BM organ to regulate its development and remodeling. Given that definitive hematopoiesis is matured in perinatal BM, it is tempting to hypothesize that the regulatory mechanisms guiding the development of bone also establish the BM niche for hematopoiesis. However, experimental evidence has been largely lacking so far. In the present study, we examined the vital role of the under-studied protein O-GlcNAcylation in determining the osteogenic versus adipogenic fate specification of BMSCs and in balancing the pro-lymphopoietic and pro-myelopoietic niche function of BMSCs. We showed that, by modifying and reciprocally regulating RUNX2 and C/EBPβ, O-GlcNAc orchestrates the early development of skeletal and hematopoietic systems (**Figure 7**).

**Figure 7.**
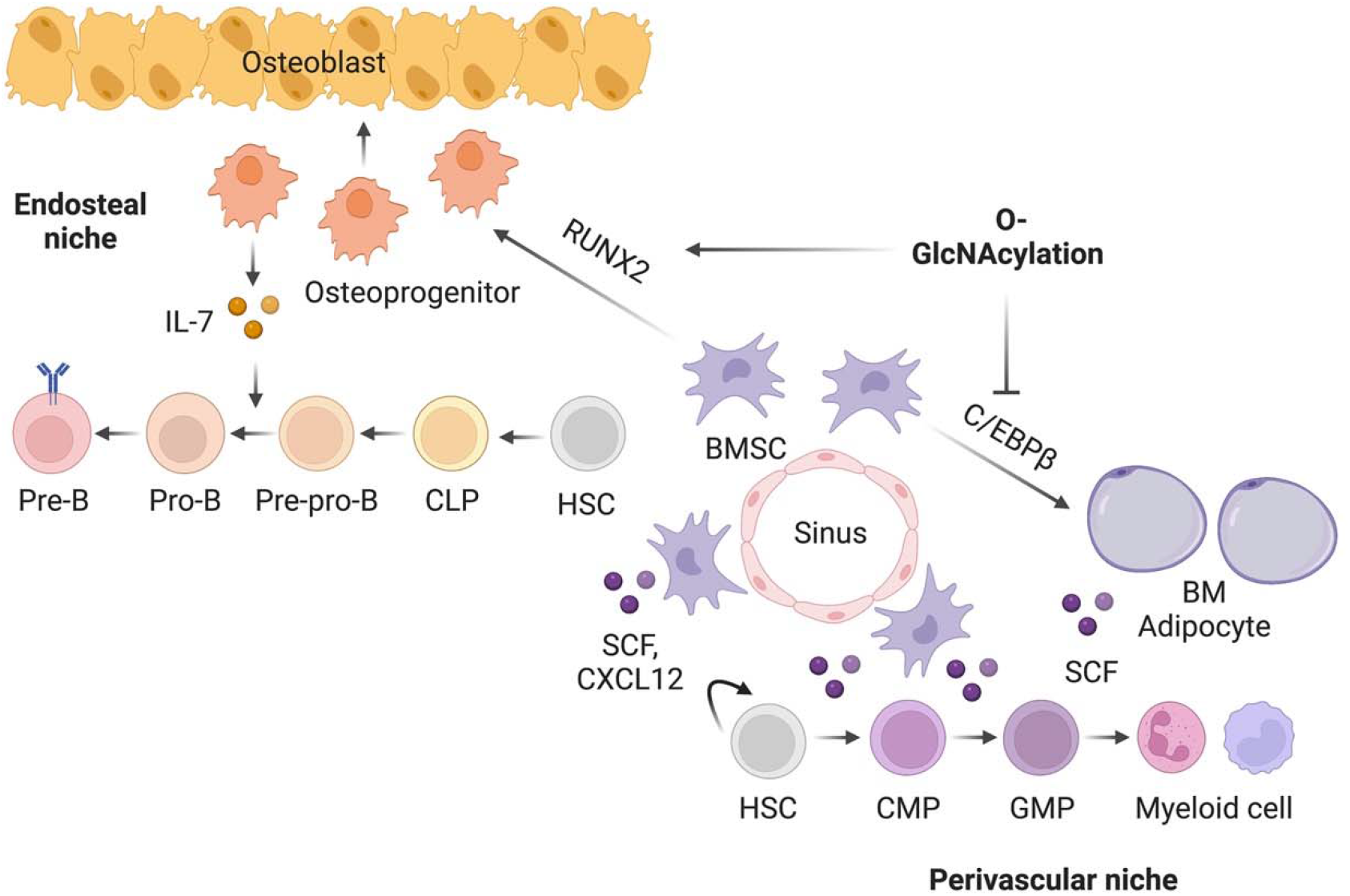
Working model of O-GlcNAc signaling in bone-BM development.

Multiple temporally and spatially distinct types of progenitors contribute to bone development and maintenance. In the early embryo, *Osx*^+^ progenitors give rise to fetal bone tissues and transient stromal cells that disappear in early postnatal life (Mizoguchi *et al*., 2014). Perinatally, *Osx*^+^ progenitors contribute to osteolineage cells and long-lived perivascular BMSCs that can be labeled by leptin receptor (*Lepr*) and adiponectin (*Adipoq*) (Zhong *et al*., 2020; Zhou *et al*., 2017). Recent evidence suggests that a significant portion of adult BMSCs and osteoblasts originate from collagen II (*Col2*)- and aggrecan (*Acan*)-expressing chondrocytes (Ono et al., 2014). Due to the fact that the *Osx-GFP:Cre* targets osteoblasts, BMSCs, and a subset of chondrocytes (Chen *et al*., 2014; Liu *et al.,* 2013), the current study could not delineate the exact developmental stages and the primary cellular compartments where OGT instructs bone development. Nonetheless, our *ex vivo* experiments and adult-onset targeting of OGT in Osx^+^ osteoblasts, together with prior published *in vivo* and *in vitro* evidence (Andres-Bergos et al., 2012; Nagel and Ball, 2014; Nagel et al., 2013), certainly reveal the indispensability of protein O-GlcNAcylation for chondro-osteogenic differentiation. While this study primarily focused early life bone development, it is warranted to further investigate the role of OGT in the transition to appositional remodeling during adulthood (Shu et al., 2021) and in osteoporosis pathogenesis during aging. Moreover, bone-forming skeletal stem cells (SSCs) are identified in other anatomical regions of long bones, such as growth plate, periosteum, and endosteum (Ambrosi et al., 2019). It remains undetermined whether O-GlcNAcylation is abundant in and controls the development and function of these SSC populations.

Protein O-GlcNAcylation senses glucose availability (Hardiville and Hart, 2014; Ruan *et al*., 2012), hormonal cues (Ruan *et al*., 2014; Ruan et al., 2017; Whelan et al., 2008), cellular stress (Martinez et al., 2017; Ruan *et al*., 2017), and immune signals (Chang et al., 2020; Liu et al., 2019; Zhao et al., 2020; Zhao et al., 2022) to maintain cellular and tissue homeostasis. Osteogenic differentiation of mesenchymal cells induces global O-GlcNAc levels (Kim *et al*., 2007; Nagel and Ball, 2014); however, the upstream mechanistic regulators of osteoblastic O-GlcNAcylation remain enigmatic. High glucose has been shown to promote O-GlcNAcylation and osteogenic differentiation of cartilage endplate stem cells (Sun et al., 2019). BMSCs preferentially use glycolysis for bioenergetics to support their self-renewal and multipotency (Ito and Suda, 2014; van Gastel and Carmeliet, 2021). Active aerobic glycolysis also fuels the high anabolic demand during bone formation. It would be important in the future to determine whether flux of the hexosamine biosynthetic pathway, a branch of glycolysis (Ruan *et al*., 2013b), increases to provide more UDP-GlcNAc for O-GlcNAc modification. We also showed here that PTH treatment increased protein O-GlcNAcylation. Signaling through the PTH receptor activates the cAMP-protein kinase A (PKA)-CREB pathway and the accumulation of inositol trisphosphate (IP3) and diacylglycerol (DAG), which further increase intracellular Ca^2+^ and PKC, respectively (Datta and Abou-Samra, 2009). Future experiments are required to determine if OGT enzymatic activity can be regulated by these signaling nodes, for example Ca^2+^/calmodulin-dependent protein kinase II (CaMKII) (Ruan *et al*., 2017).

The BM microenvironment, composed of BMSCs, osteoblasts, adipocytes, sympathetic nerves, and vascular endothelial cells, has been highlighted as an important extrinsic factor for the maintenance and differentiation of distinct hematopoietic lineage progenitors (Bianco and Robey, 2015; Calvi and Link, 2015; Morrison and Scadden, 2014; Wei and Frenette, 2018). While the concomitant development, remodeling, and aging of the skeletal and hematopoietic systems have been observed in various pathophysiological conditions, mechanisms underlying the coordinated regulation of the two systems are less understood. Our current study has provided the first evidence that RUNX2, permitted by O-GlcNAcylation, not only is indispensable for the osteoblast development, but also establishes the endosteal niche for B lymphocytes by driving IL-7 expression (**Figure 7**). When OGT is deficient, the perivascular BMSCs are prone to adipogenic differentiation, which also activates C/EBPβ-dependent SCF expression and myelopoiesis. During aging, the parallel dysfunction of the skeletal and hematopoietic systems leads to osteoporosis, marrow fat accumulation, and myeloid hematopoietic skewing (Geiger et al., 2013). Whether BMSC aging is associated with O-GlcNAc decline and whether the balance between RUNX and C/EBPβ leads to bone-fat imbalances and niche dysfunction require future investigations.

## METHODS

### Animals

All animal experiments were approved by the institutional animal care and use committee of the University of Minnesota. All the mice were group-housed in light/dark cycle- (6am-8pm light), temperature- (21.5 ± 1.5 ^o^C), and humidity-controlled (30-70%) room, and had free access to water and regular chow (Teklad #2018) unless otherwise indicated. All mice were maintained on a C57BL6 background. Due to the X-chromosome localization of the *Ogt* gene, only male mice were used in the study if not specified in the text or figures. To suppress Cre activity, designated breeders were fed a diet containing 200 mg/kg doxycycline (Bio-serv, S3888).

### BMSC isolation, culture, and differentiation

BMSC were isolated from the long bones as described previously (Zhu et al., 2010). The fragments of long bones were digested with collagenase II for 30 mins. The released cells were discarded, and the digested bone fragments were cultivated in the BMSCs growth medium (alpha-MEM supplemented with 10% FBS). Once confluent, cells were switched to either adipogenic differentiation medium (alpha-MEM supplemented with 20% FBS, 500 μM IBMX, 1 μM Dexamethasone, 10 μg/ml Insulin and 1 μM Rosiglitazone) for the first 2 days. The medium was then changed to adipocyte differentiation base medium (α-MEM supplemented with 20% FBS, 10 μg/ml Insulin and 1 μM Rosiglitazone) for the next 4 days followed by oil red O staining. For osteogenic differentiation, cells were induced with osteoblast differentiation medium (α-MEM supplemented with 10% FBS, 0.3mM ascorbic acid, 10 mM β-glycerophosphate, 0.1 μM Dexamethasone) for 14 days followed by ALP staining or for 28 days followed by Alizarin red staining.

### Cell culture, plasmids, and lentiviruses

HEK 293, COS7, and C3H10T1/2 (ATCC, CCL-226) cells were cultured with DMEM plus 10% of FBS. The mouse RUNX2-Myc/DDK plasmid was purchased from OriGene (MR227321), then subcloned into pLV-EF1a-IRES-Hygro (Addgene #85134). Mouse PPARλ2 with a N-terminal MYC tag was subcloned into pLVX-Dsred-puro plasmid. C/EBPβ plasmids were kindly provided by Dr. Xiaoyong Yang at Yale University and then subcloned into pCDH-CMV-P2a-Puro. O-GlcNAc site were mutated into alanine with Q5^®^ Site-Directed Mutagenesis Kit (NEB#E0554). Lentivirus were packed as previously described (Huang et al., 2022). Briefly, 293FT cells were transfected with over-expression plasmids and pSPAX2 / pMD2G. Mediums with lentivirus were filtered and added into C3H10T1/2 cells. 72h after infection, cells were selected with drugs according to the resistance genes they possessed.

### Luciferase assay

For Runx2 luciferase assay, with empty or RUNX2 vectors were transfected into COS7 cells with Lipofectamine, together with 6xOSE2-luc and pGL4-hRluc vectors in which either firefly or Renilla luciferase genes were expressed under the control of the RUNX2-specific or the constitutive SV40 promoter, respectively. After 6 h cells were washed three times and with the addition of 50 uM OSMI-1. Cells were incubated for an additional 48 h in growth medium containing 5% serum. Luminescent signals were generated using the Dual-Luciferase Assay System (Promega). Relative light units (RLU) for the 6xOSE2 reporter were normalized against pGL4-hRLuc values as an internal control for transfection efficiency. For PPARγ2 luciferase assays, C3H10T1/2 cells were transfected with Transporter 5^®^ Transfection Reagent (Polysciences) following manufacture’s protocol. PPARγ2 transcriptional activity was determined using the PPREx3-TK-luc reporter (Addgene, #1015).

### Histology

Bone tissues were fixed in formalin solution at 4 ^o^C for 24 h. Tissue embedding, sectioning, and hematoxylin and eosin staining were performed at the Comparative Pathology Shared Resource of the University of Minnesota. For immunostaining, the tissues were embedded in OCT then cut into 7-μm slides. After three times of PBS wash, the slides were incubated with blocking buffer (3% BSA in PBS) for 1 h, then immersed with anti-Perilipin (Cell Signaling Technology, #9349) antibody overnight at 4°C. For immunofluorescence, PBS-washed slides were incubated with a fluorescent secondary antibody at room temperature for 1 h, and then mounted with VECTASHIELD^®^ Antifade Mounting Medium with DAPI after three times of PBS wash. A Nikon system was used for imaging. Goldner’s trichrome and Safranin O staining were performed at Servicebio, China.

### micro-CT

The samples were scanned with an in vitro micro-CT device (Skyscan 1272, Bruker micro-CT) with scanning parameters of: Source Voltage = 60 kV, Source Current = 166uA, exposure 897 ms/frame, average of 3 frames per projection, Rotation Step (deg) = 0.200- and 0.25-mm Aluminum filter. The specimens were scanned at high resolution (2016 × 1344 pixels) with an Isotropic voxel size of 7.1 μm. Reconstructions for X-ray projections and re-alignment were performed using the Skyscan software (NRecon and DataViewer) (v. 1.7.3.1, Brüker micro-CT, Kontich, Belgium). Ring artefact and beam hardening corrections were applied in reconstruction. Datasets were loaded into SkyScan CT-Analyzer software for measurement of BMD. Calibration was performed with 0.25- and 0.75-mg/mL hydroxyapatite mice phantoms provided by SkyScan. For cancellous and cortical bone analysis, the scanning regions were confined to the distal meta-physis, 100 slices starting at 0.5 mm proximally from the proximal tip of the primary spongiosa for the cancellous portion and 100 slices starting at 4.5 mm proximally from the center of intercondylar fossa for the cortical portion.

### O-GlcNAc mass spectrometry

Myc-tagged PPARγ2 was co-transfected with OGT into 15 cm-dishes of 293T cells and purified by immunoprecipitation with anti-c-Myc agarose beads (Pierce), followed by PAGE gel electrophoresis. The corresponding PPARγ2 band was cut for in gel Trypsin (Promega) digestion. Tryptic peptides were analyzed by on-line LC-MS/MS using an Orbitrap Fusion Lumos (Thermo) coupled with a NanoAcquity UPLC system (Waters) as we previously reported (Liu *et al*., 2019; Zhao *et al*., 2022). Peaklists were generated using PAVA (UCSF) and searched using Protein Prospector 5.23.0 against the SwissProt database and a randomized concatenated database with the addition of the recombinant PPARγ2 sequence. HexNAcylated peptides were manually verified.

### Real-time RT-PCR

RNA was isolated with Trizol and reverse transcribed into cDNA with the iScript^™^ cDNA Synthesis Kit. Real-time RT-PCR was performed using iTaq^™^ Universal SYBR^®^ Green Supermix and gene-specific primers (**Figure 6 —source data 1**) on a Bio-Rad C1000 Thermal Cycler.

### Flow cytometry

For B-cell lymphopoiesis, BM cells were stained in PBS containing 1% (w/v) bovine serum albumin on ice for 30 min, with anti-B220 (Life Technologies, 67-0452-82), anti-CD43 (BD Biosciences, 553271), anti-CD24 (Biolegend, 101822), anti-Ly-51 (Biolegend, 108305), anti-CD127 (Tonbo Biosciences, 20-1271-U100), anti-CD25 (Life Technologies, 63-0251-82) and anti-CD19 (Biolegend, 115545). For hematopoietic stem and progenitor cells, BM cells were stained with a cocktail of biotin-conjugated lineage antibodies CD3e, B220, Ter119, Mac-1 and Gr-1 (Biolegend, 133307), CD4 (Biolegend, 100403), CD5 (Biolegend, 100603), CD8 (Biolegend, 100703), followed by Streptavidin-AF488 (Biolegend, 405235). Cells were then stained with CD127-APC (eBioscience, 17-1271-82), c-Kit-APC-eFluor780 (eBioscience, 47-1171-82), Sca-1-Super Bright 436 (eBioscience, 62-5981-82), CD34-PE (Biolegend, 152204) and FcγR-PerCP-eFluor710 (eBioscience, 46-0161-80), CD150-BV605 (Biolegend, 115927), and CD48-BUV395 (BDBioscience, 740236). Fixable Viability Dye was used to exclude dead cells as instructed by the manufacturer. A complete list of used antibodies was shown in **Figure 5 —source data 1**. Flow cytometry was performed on an LSR Fortessa H0081 or X20 and analyzed with FlowJo.

### Quantification and statistical analysis

Results are shown as mean ± SEM. N values (biological replicates) and statistical analysis methods are described in figure legends. The statistical comparisons were carried out using two-tailed unpaired Student’s t-test and one-way or two-way ANOVA with indicated post hoc tests with Prism 9 (Graphpad). Differences were considered significant when p < 0.05. *, p < 0.05; **, p < 0.01; ***, p < 0.001.

## ACKNOWLEGEMENT

We thank Dr. Xiaoyong Yang for kindly providing wildtype and mutant C/EBPβ plasmids. This work was supported by the National Natural Science Foundation of China, China (32170847) to Z.H., NIH/NIAID (R01 AI162678) to H.Z., and NIH/NIAID (R01 AI139420 and R01 AI162791) to H.-B.R.

## DATA AVAILABILITY

All data generated and analyzed are included in the manuscript. Source data files are provided for relevant figures.

## AUTHOR CONTRIBUTION

Z.Z. and Z.H. designed and performed experiments, analyzed data, and contributed to manuscript writing. M.A., M.E., J.C., and K.C.M. performed micro-CT scanning and analysis. L.E.B. determined RUNX2 O-GlcNAcylation and transcriptional activity. H.Z. provided guidance and assistance on flow cytometry. J.C.M. and A.L.B. performed mass spectrometric identification of PPARγ2 O-GlcNAc sites. H.-B.R. conceived the project, designed experiments, analyzed data, and wrote the manuscript.

## DECLARATION OF INTERESTS

The authors declare no competing interests.

## SOURCE DATA LEGNED

**Figure 2 —source data 1**

Raw uncropped image for panel D.

**Figure 2 —source data 2**

Raw uncropped image for panel G.

**Figure 4 —source data 1**

Mass spectrometry Peaklists of all protein modifications (Table S1) and PPARγ2 O-GlcNAc sites (Table S2).

**Figure 5 —source data 1**

Antibodies used for flow cytometry.

**Figure 6 —source data 1**

Sequences of oligos used for RT-qPCR.

## SUPPLIMENTARY FIGURES

**Figure 1—figure supplement 1.**
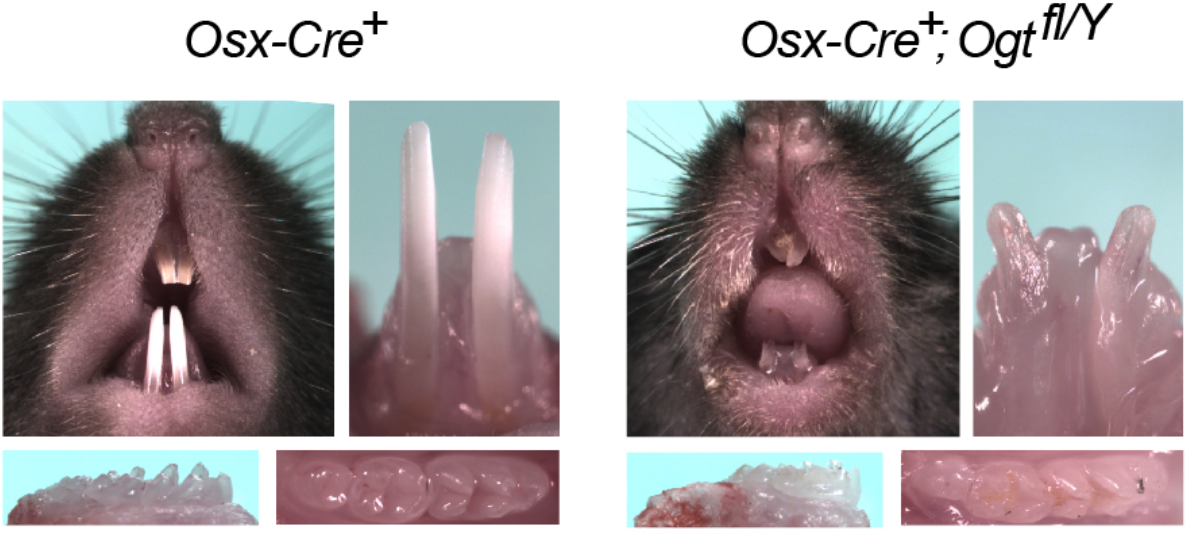
Dental defects of *Osx*^Δ*Ogt*^ mice. Top: photographic analysis of incisor tooth development. Bottom: photographic analysis of molar tooth development.

**Figure 4—figure supplement 1.**
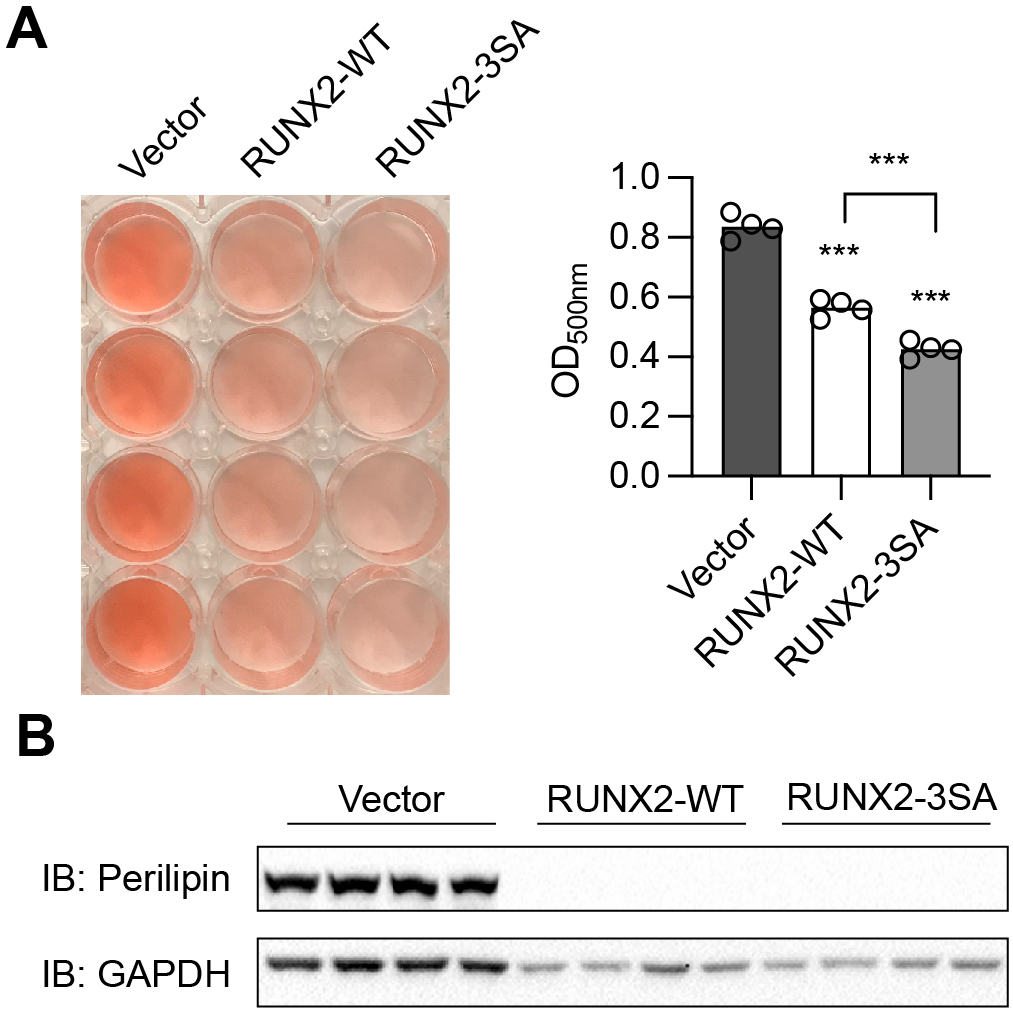
RUNX2 inhibits adipogenesis independently of O-GlcNAcylation. C3H10T1/2 cells were infected with lentiviruses expressing WT- or O-GlcNAc-deficient (3SA)-RUNX2, then induced for adipogenic differentiation. Adipogenesis was quantified by Oil Red O staining (**A**) and Perilipin Western blotting (**B**). Data are presented as mean ± SEM.***, p < 0.01 by one-way ANOVA.

**Figure 4—figure supplement 2.**
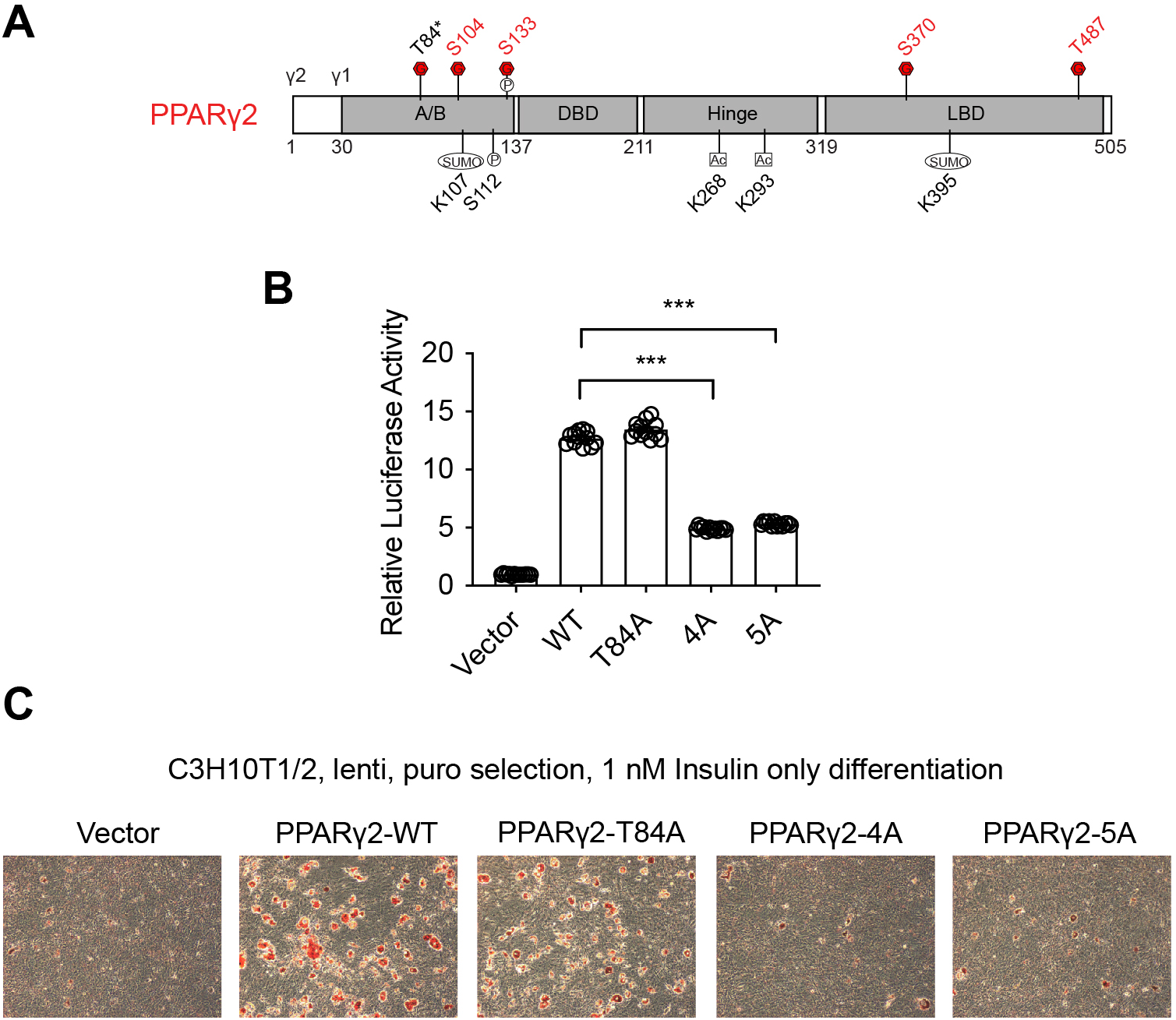
PPARγ2 O-GlcNAcylation is required for adipogenesis. (**A**) PPARγ2 protein domains with the O-GlcNAc site shown in red. (P)hosphorylation, (SUMO)lation and (Ac)etylation sites were also shown. (**B**) Previously known (T84) and newly identified (S104, S133, S370, and T487) O-GlcNAc sites were mutated to alanine separately (T84A and 4A, respectively) or together (5A). PPRE-luciferase assay of PPARγ2 was performed. Data are presented as mean ± SEM.***, p < 0.01 by one-way ANOVA. (**C**) C3H10T1/2 cells were infected with lentiviral PPARγ2, induced to become adipocytes in the presence of only insulin, and stain with Oil Red O.

**Figure 5—figure supplement 1.**
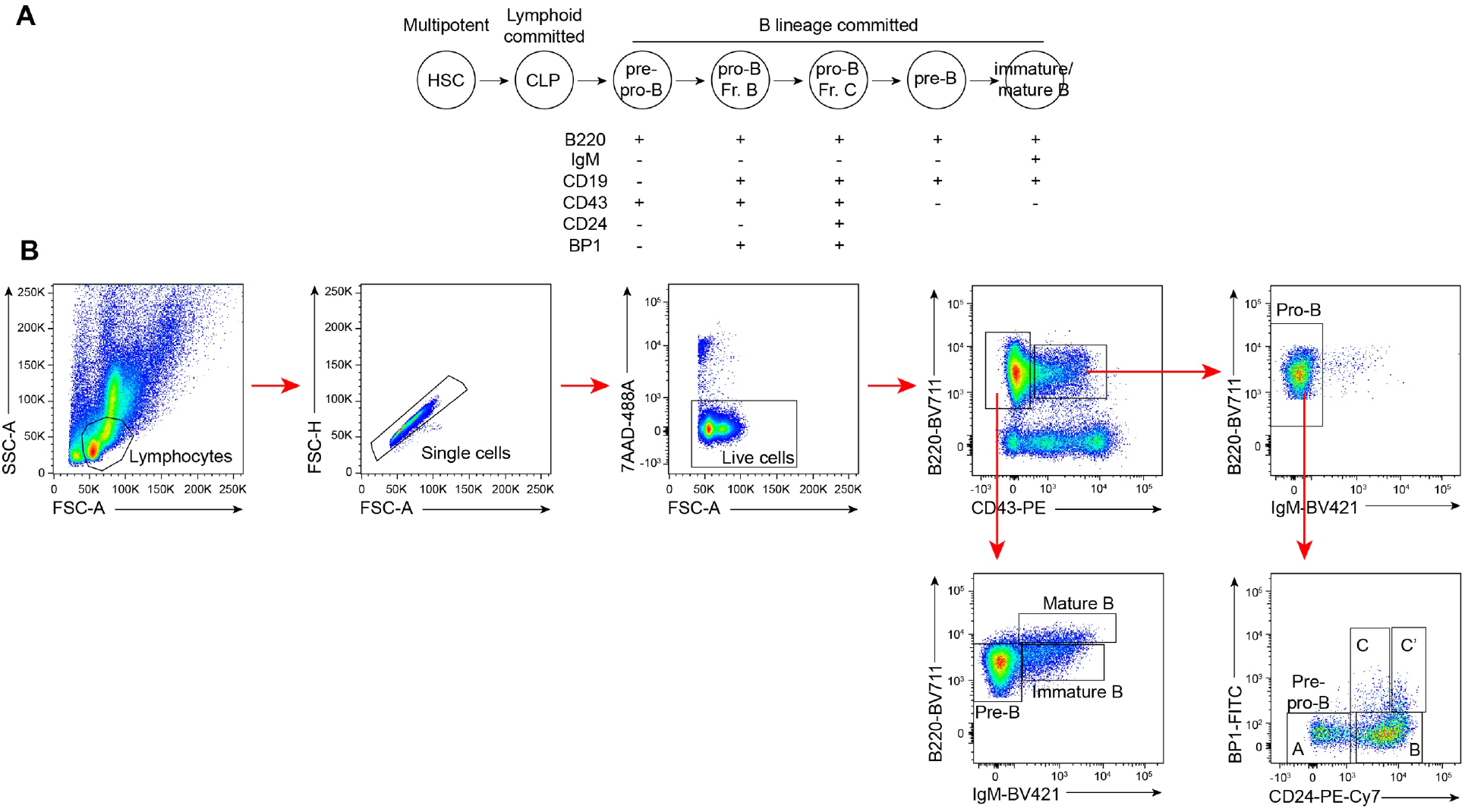
Flow cytometry of BM B cells. (**A**) B cell lineage development and surface markers used for flow cytometry. (**B**) Gating strategy for B cell populations.

## Notes

### Competing Interest Statement

The authors have declared no competing interest.

